# Metagenomics insights into microbial diversity shifts among seagrass sediments

**DOI:** 10.1101/2024.01.14.574613

**Authors:** Matsapume Detcharoen, Ekkalak Rattanachot, Anchana Prathep

## Abstract

Seagrasses are important in marine ecosystems by providing food and shelter for animals, storing carbon, and reducing sulfide and methane emissions. Previous studies found variations in the microbiomes of seagrass both above and below sediment surface, with certain microbes serving as indicators of seagrass health. Despite numerous studies on bacterial communities in specific seagrass species, little is known about bacterial communities in mixed seagrass beds. This study used metagenomics to investigate microbial diversity in bare and seagrass-associated sediments, across seven seagrass species combinations. The findings revealed higher alpha diversity of bacteria and archaea in the upper layers, whereas microbial eukaryotes were more diverse in the lower layers. Significant taxonomic differences were found between bare and seagrass sediments, and among seagrass species. Several metagenome-assembled genomes were reconstructed, primarily from Pseudomonadota and Thermodesulfobacteriota. This study improves our understanding of complex interactions between seagrasses and their associated microbial communities and laying a foundation for efficient strategies for the management and restoration of seagrass ecosystems.

## Introduction

Seagrasses are marine flowering plants with a significant ecological role in marine ecosystems. They are food sources for herbivores and provide essential habitats for numerous marine animals. Seagrasses are the primary carbon source in the detrital pool of the marine environment, facilitating carbon storage (Stankovic et al., 2021) and reducing sulfide and methane emissions (Lyimo et al., 2018). They bury organic carbon in the seabed faster than terrestrial ecosystems (McLeod et al., 2011). Their leaves and roots can filter and trap nutrients and bacteria in water, and modifying wave patterns (Lamb et al., 2017; Liu et al., 2023). Consequently, seagrasses are crucial in maintaining the health and equilibrium of coastal ecosystems by offering shelter, sustenance, and nursery grounds for various organisms, including fishes, invertebrates, and certain marine mammals.

Recently, the microbial diversity of seagrass has become a research focus using traditional cultivation and next-generation sequencing methods. The associated microorganisms are essential for nutrient cycling (e.g., nitrogen fixation), carbon storage (Mou et al., 2008), and seagrass growth (Celdrán et al., 2012). Additionally, microbes, such as methylotrophic and iron-cycling bacteria, found near seagrass roots are good indicators of the health status of seagrass beds (Martin et al., 2020). Although numerous bacterial communities in specific seagrass species, such as *Halophila ovalis* (Martin et al., 2020; Titioatchasai et al., 2023), *Thalassia hemprichii* (Jiang et al., 2015a), and *Zostera muelleri* (Brodersen et al., 2018) have been investigated, the bacterial communities in mixed seagrass beds are poorly understood.

Despite harboring 13 seagrass species (Fortes et al., 2018), the microbial diversity of seagrass beds in Thailand is relatively unknown. This study aimed to discover the microbial diversity in sediments associated with seven combinations of seagrass species and bare sand sediments in the same area using metagenomics. Our findings revealed significant differences in several taxa between seagrass and bare sand sediments. Our findings contribute to the knowledge about microbial diversity in seagrass, revealing the potential of these ecosystems to combat climate change.

## Methods

### Sample collection, DNA extraction, and sequencing

Sediments were collected using core samples (7 cm in diameter and 5 cm depth) in intertidal areas at Mook island, Trang, Thailand (7°22’42.68”N, 99°18’27.18”E), on October 23, 2021. All samples were stored on ice until they arrived at the laboratory on the same day. The samples consisted of Bare Sands (BS) and seven seagrass sediments with over 70% coverage, including mono-patches of *Enhalus acoroides* (Ea), *H. ovalis* (Ho), *T. hemprichii* (Th) and mixed patches of *T. hemprichii* and *E. acoroides* (Th+Ea), *T. hemprichii* and *H. ovalis* (Th+Ho), *H. ovalis* and *E. acoroides* (Ho+Ea), and *T. hemprichii*, *H. ovalis*, and *E*. *acoroides* (Th+Ho+Ea). Three samples were taken for each sediment type. Because the mean root length for Ea, Ho, and Th were 7.7, 6.5, and 2.1 cm, respectively (Duarte et al., 1998), we divided each sediment into two parts: the upper layer (0–2 cm depth) and the lower layer (4–5 cm depth).

DNA extraction was done using the DNeasy PowerSoil Pro kit (QIAGEN, Hilden, Germany), following the manufacturer’s protocol. DNA quality was checked using 1% agarose gel electrophoresis. Two out of the 48 samples were discarded because of an insufficient amount of DNA, resulting in a total of 46 samples. Whole-genome metagenomic 2 × 150-bp paired-end sequencing using BGISEQ was performed at BGI Genomics (Hong Kong, China). Raw sequence quality was checked, and adaptors were trimmed using fastp v0.23.0 (Chen et al., 2018) with default parameters. The raw sequence data generated in this study were deposited in the NCBI Sequence Read Archive under the project number PRJNA862559.

### Taxonomy assignment and differential abundances

The reads of each sample were taxonomically classified using the Kaiju Web server (Menzel et al., 2016). The non-redundant protein database (NCBI BLAST nr+euk v2021-03), which includes bacteria, archaea, fungi, and microbial eukaryotes, was used as the reference database. Several parameters were applied, including filtering out low complexity, running in greedy mode, setting a minimum match length of 11, minimum match score of 75, maximum mismatch of five, and maximum E-value of 0.01, and otherwise default settings. Sequences with unassigned taxa were excluded from this dataset.

We performed microbial diversity analysis using phyloseq v1.40.0 (McMurdie and Holmes, 2013). To measure the alpha diversity, all samples were rarefied to the minimum reads found among all samples, and the Simpson index was calculated. Significant differences in alpha diversity were tested using the Kruskal-Wallis test, with an alpha of 0.05. Non-metric multidimensional scaling (NMDS) analysis and analysis of similarities (ANOSIM) were performed based on the Bray-Curtis distance for beta diversity. Differential abundances of microbial taxa were determined using linear discriminant analysis effect size (LEfSe) v1.1.01 (Segata et al., 2011).

For functional annotation, the metagenomic reads of each sample were assembled into contigs using metaSPAdes v3.15.3 (Prjibelski et al., 2020). The genes were annotated using Prokka v1.14.5 (Seemann, 2014) with default options. The amino acid sequences obtained from this process were used as queries for GhostKOALA (Kanehisa et al., 2016) to obtain the Kyoto Encyclopedia of Genes and Genomes (KEGG) pathways.

### Metagenome assembled genomes analysis

We used the KBase platform (Arkin et al., 2018) for read assembly, contig binning, and assessing the quality of metagenome-assembled genomes (MAGs). We pooled reads from the samples of the same sediment and assembled into contigs using metaSPAdes function in SPAdes v3.15.3 (Prjibelski et al., 2020), with the optimal k-mer parameter and a minimum contig length of 1000 bp. Assembly statistics were calculated using QUAST v5.0.2 (Mikheenko et al., 2018). Contig binning was performed using MaxBin v2.2.4 (Wu et al., 2016), CONCOCT v1.1 (Alneberg et al., 2014), and MetaBat2 v2.3.0 (Kang et al., 2019) with default parameters. The DAS Tool v1.1.2 (Sieber et al., 2018) was used to combine bins from the different binning programs in the previous step. The quality of the binned genomes was assessed using CheckM v1.0.18 (Parks *et al*., 2015), and genomes with less than 75% completeness and more than five percent contamination were discarded. We used DRAM v1.2.4 (Shaffer et al., 2020) to annotate the genomes, with a minimum contig length of 500 and a bit score threshold of 60. The taxonomic classification and phylogenetic relationships of the MAGs were determined using GTDB-Tk v2.3.2 with the reference data version r214 (Chaumeil et al., 2020).

## Results

### Microbial diversity

Of all 46 samples, the mean number of reads per sample was 7,099,696. In total, 60,634 operational taxonomic units were identified. There were 3,282 viruses, 48,483 bacteria, 2,620 archaea, and 6,249 microbial eukaryotes. The sediment from Ea had the most bacterial reads, whereas Ho+Ea had the most archaeal reads (Figure 1A, Table S1). The mixed patches of Th+Ho+Ea had the most eukaryotic reads. Contrastingly, Th had the fewest bacterial and microbial eukaryotic reads, whereas Th+Ho+Ea had the lowest number of Archaeal reads.

**Figure 1.**
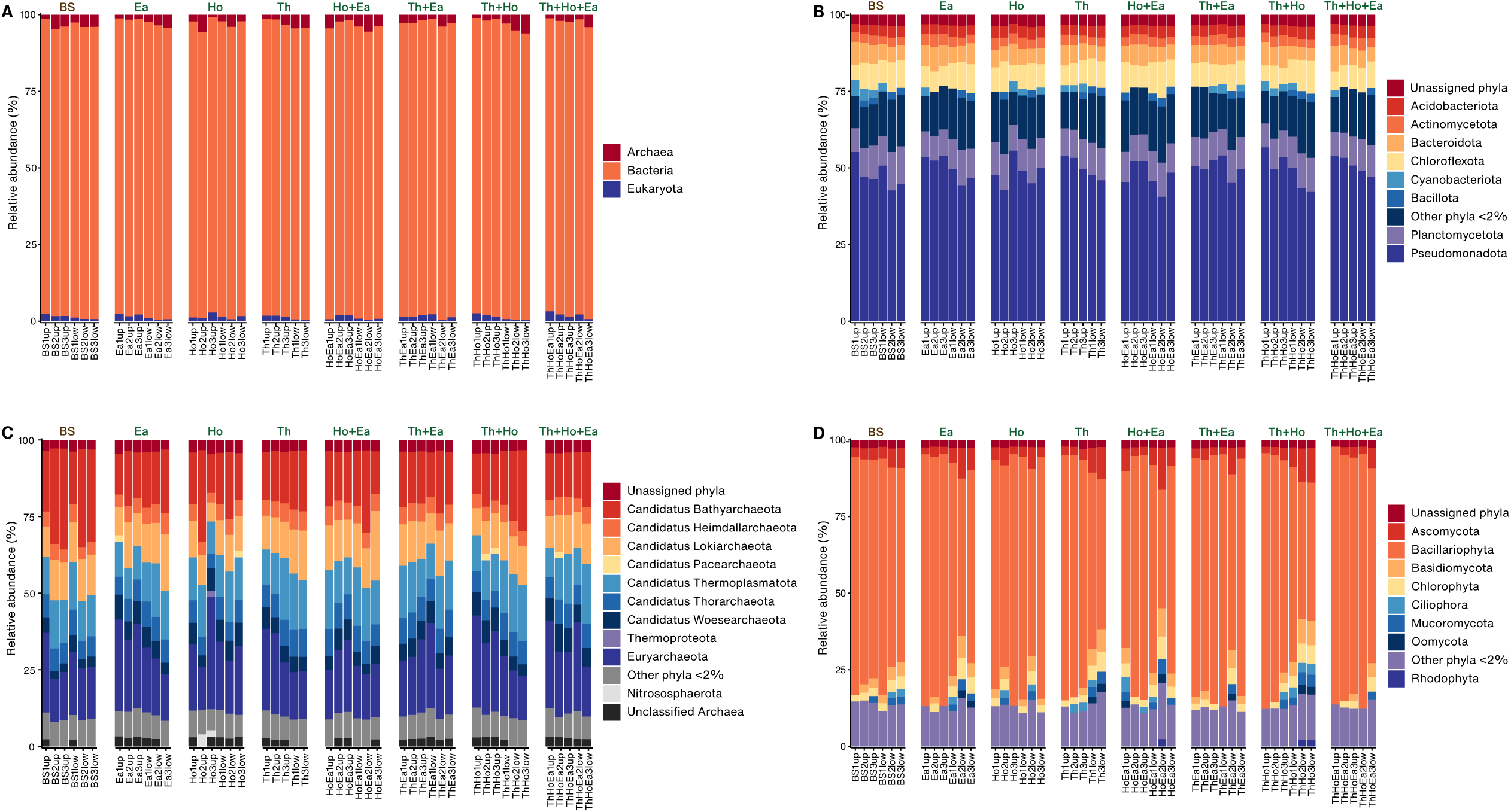
Bar plots represent relative abundance of cellular organisms in each sample. At the superkingdom level (A), the phylum level of bacteria (B), archaea (C), and microbial eukaryotes (D).

Among the bacterial phyla, Pseudomonadota (Proteobacteria) was the most abundant in all samples, followed by Planctomycetota (Planctomycetes) and Chloroflexota (Chloroflexi) (Figure 1B). Among the archaea, the most prevalent phyla were *Candidatus* Bathyarchaeota, Euryarchaeota, and *Candidatus* Thermoplasmatota (Figure 1C). Additionally, Bacillariophyta was the most abundant microbial eukaryotic phylum in all samples (Figure 1D).

### Upper and lower sediment layers

Comparisons of the upper and lower layers of the sediments revealed that Pseudomonadota was the predominant bacterial taxon in both the upper and lower layers, whereas Planctomycetota was more abundant in the lower layers. Among the archaea, *Ca*. Bathyarchaeota, *Ca*. Thermoplasmatota, and *Candidatus* Lokiarchaeota were predominant in the lower layers, whereas Euryarchaeota dominated in the upper layers. In eukaryotes, Bacillariophyta (i.e., diatoms) was the predominant taxon in the upper layers, whereas Chlorophyta and Basidiomycota were the two most abundant taxa in the lower layers (Figure S1, Figure S2).

Overall, the alpha diversity was significantly higher in the upper layer than in the lower layer (Kruskal-Wallis p = 0.0001, Table S2). Bacterial and archaeal diversity in the upper part of the sediment was higher than that in the lower part (p = 0.0001 and 0.0046 respectively), whereas microbial eukaryotes showed the opposite pattern (p = 0.0007).

When each sediment was considered separately, the upper layers of BS and Ho differed from their respective lower layers (Kruskal-Wallis p = 0.0495 for both, Figure 2A). The bacterial alpha diversity in the upper layers of BS and Th+Ho was higher than in their lower layers (p = 0.0495 for both, Figure 2B). Furthermore, the upper layers of both Ea and Th+Ho exhibited greater archaeal diversity than their respective lower layers (p = 0.0495 for both, Figure 2C). Microbial eukaryotes were more diverse in the lower layer than the upper layer, with significant values for Ea and Th+Ho (p = 0.0495 for both, Figure 2D).

**Figure 2.**
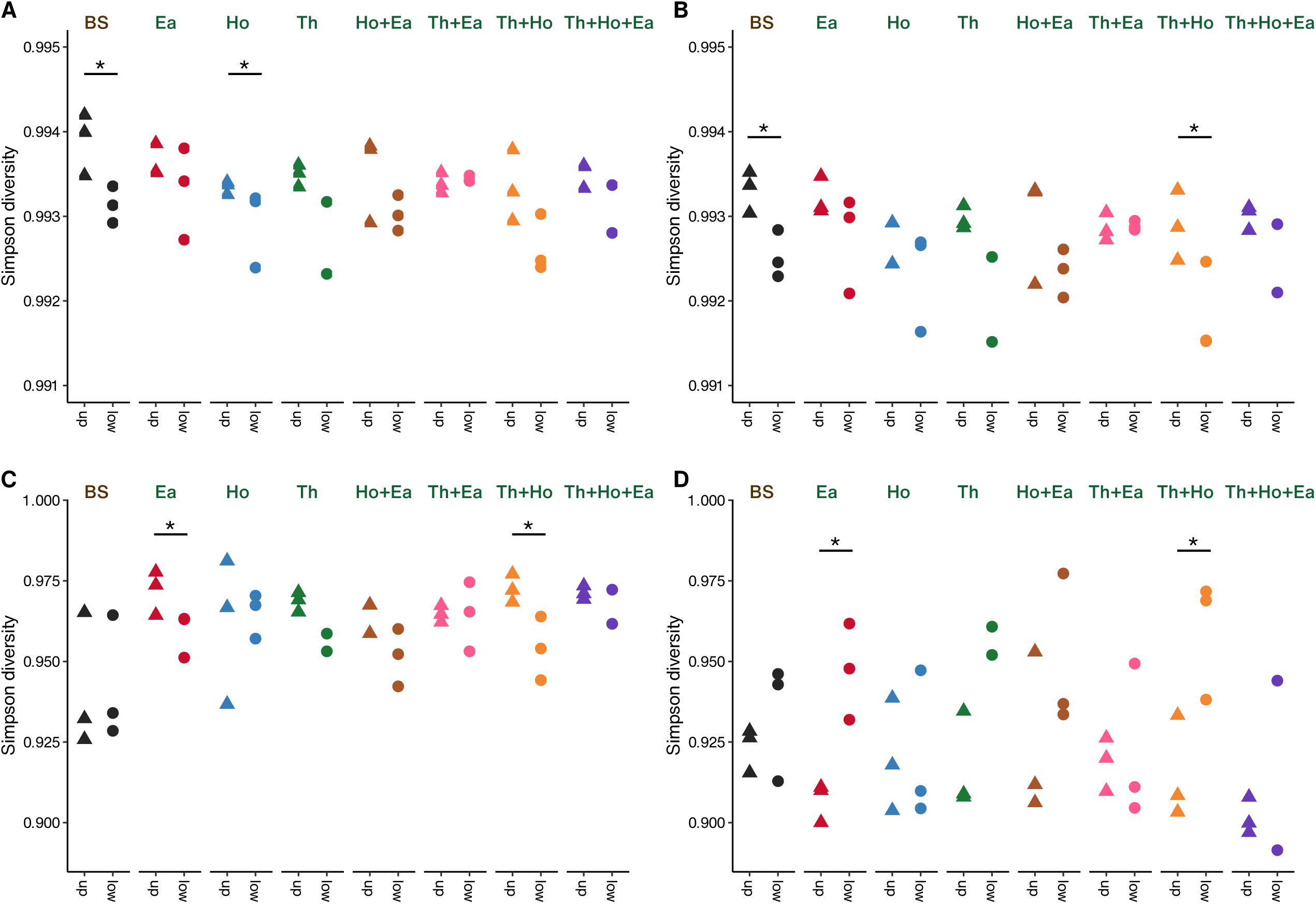
Alpha diversity based on the Simpson index between the upper and lower layers of Bare Sand (BS) and seagrass-covered sand. All the taxa included (A), bacteria (B), archaea (C), and eukaryotes (D).

Beta diversity, assessed using NMDS, revealed slight dissimilarity between the upper and lower layers. The upper and lower layers of Th+Ho, Ea, and Th exhibited the greatest dissimilarity, whereas Ho, Th+Ea, Ho+Ea, and BS showed similarity (Figure 3A, Figure S3). The highest degree of similarity was observed when only eukaryotic organisms were analyzed (ANOSIM statistic R = –0.09172, Figure 3D).

**Figure 3.**
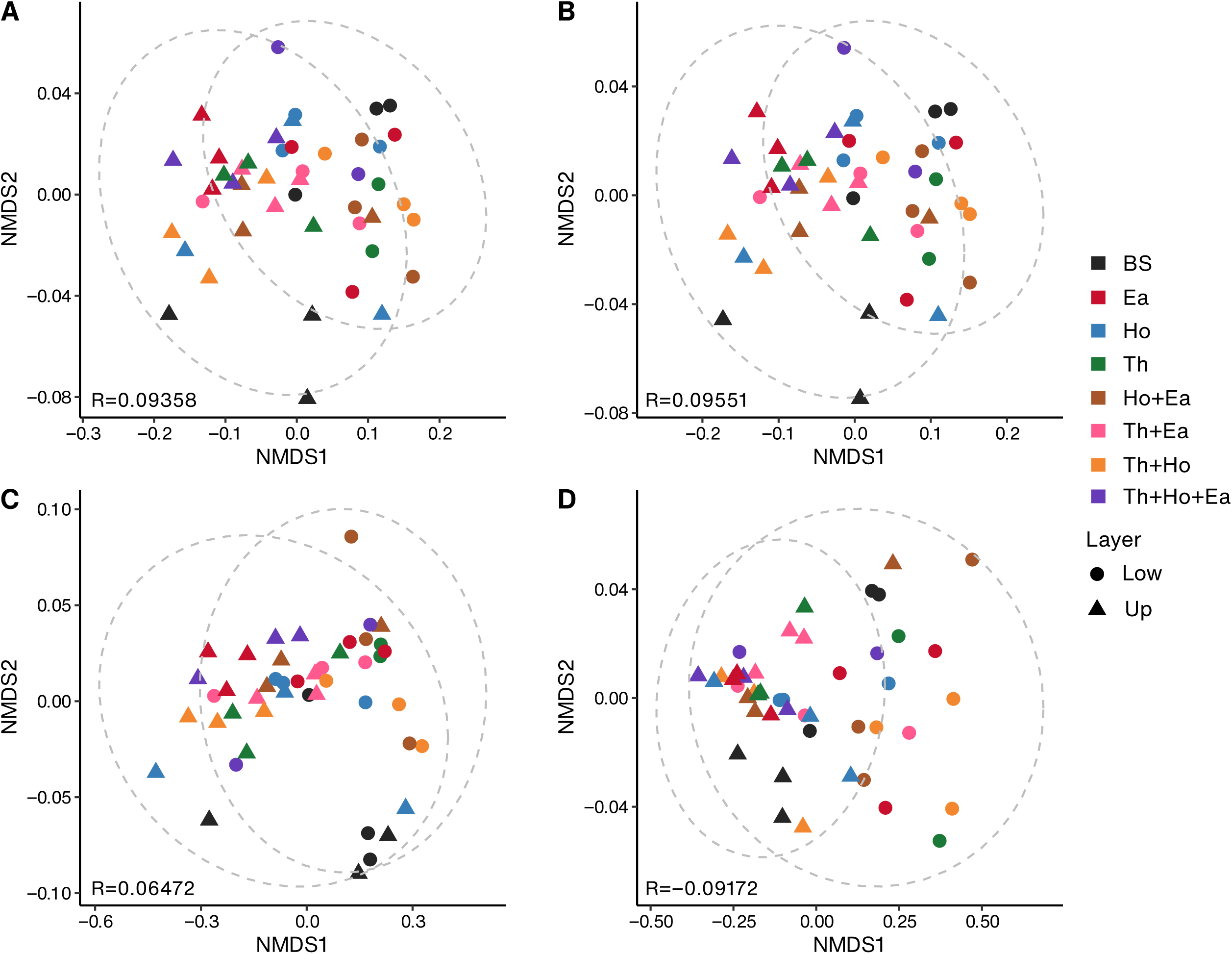
Nonmetric multidimensional scaling based on the Bray-Curtis distance of all samples. All the taxa included (A), bacteria (B), archaea (C), and microbial eukaryotes (D). ANOSIM statistic R values are shown at the bottom left.

Within the upper layers, sulfur-fixing bacteria, such as *Thiohalobacter*, *Thiogranum*, and *Thiohalophilus*, were enriched in Th+Ho+Ea, while nitrogen-fixing cyanobacteria, such as *Crocosphaera* and *Rippkaea*, were more prevalent in BS (Figure 4A). In the lower layers, seagrass-associated sediments exhibited a higher significant abundance of taxa, mainly chemoheterotrophic bacteria such as *Haliangium*, *Desulfatitalea*, and *Rhodopirellula.* In contrast, *Sulfurovum* sp. was the only taxon enriched in the lower layer of BS (Figure 4B).

**Figure 4.**
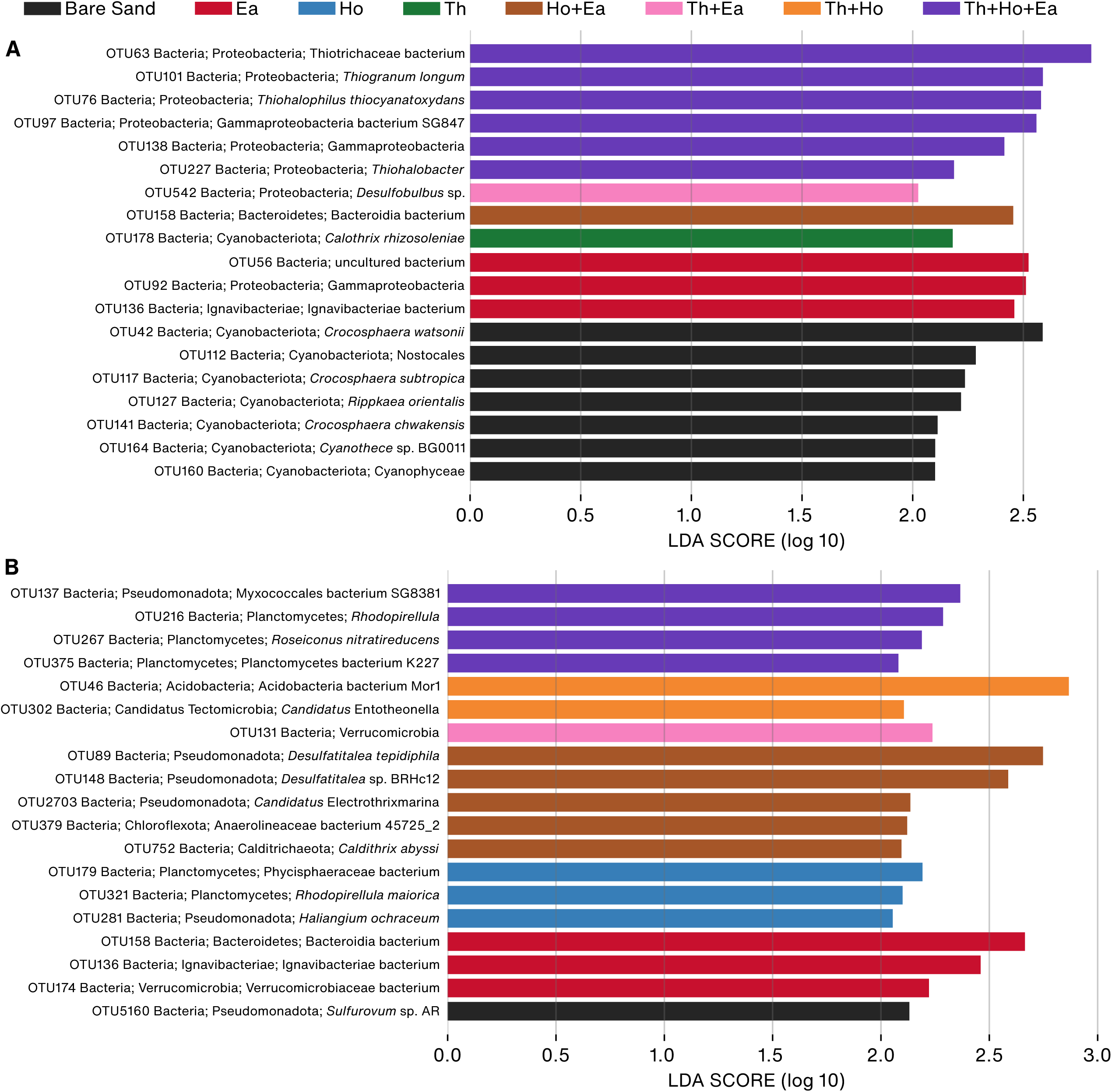
Linear discriminant analysis effect size (LEfSe) showed significantly enriched taxa in the upper (A) and lower (B) layers of the sediments.

### Bare Sand and seagrass

We compared the alpha diversity of microbial communities in the seagrass-associated sediments and BS. Among all taxa, the mean alpha diversity of all seagrass samples was 0.993, and 0.994 for BS (Table S2). There was no significant difference in alpha diversity between the BS and seagrass samples (Kruskal-Wallis test, p = 0.4148). Beta diversity, a comparison between BS and seagrass soils, showed high similarity (ANOSIM statistic R = 0.09358, significance = 0.117).

We identified seven distinct bacterial taxa in BS and seagrass using differential abundance analysis. Interestingly, several Pseudomonadota taxa were significantly enriched in seagrass, whereas several cyanobacteria taxa were more abundant in BS (Figure 5A).

**Figure 5.**
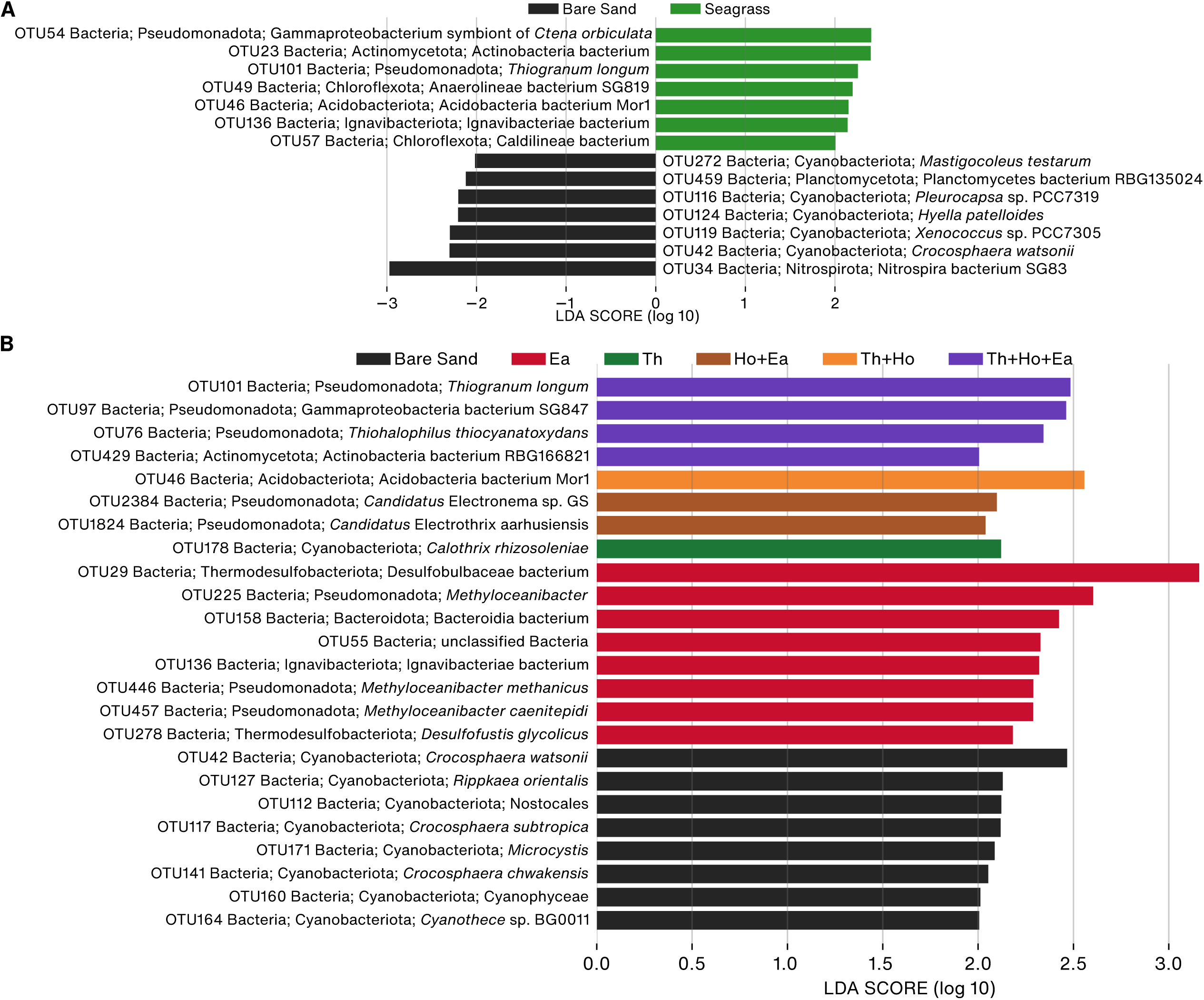
Differential abundance analysis using linear discriminant analysis effect size (LEfSe) calculated performed for all taxa. Consider Bare Sand and all seagrass samples (A), and all different types of Bare Sand and seagrass samples (B).

When all samples and the upper and lower layers were analyzed together, several cyanobacterial taxa were significantly abundant in the BS (Figure 5B). Ea exhibited the highest number of significantly abundant taxa, including facultative methylotrophic bacteria *Methyloceanibacter*. Furthermore, sulfur-reducing bacteria, such as *Thiohalophilus* and *Thiogranum*, and those belonging to Desulfobulbaceae were enriched in Ea and Th+Ho+Ea.

### Differences between Bare Sand and different seagrass species

Comparative analysis between BS and various types of seagrass sediments indicated a higher diversity of taxa in seagrass sediments than in BS. Notably, seagrass patches consisting of Ea (Ea, Ho+Ea, Th+Ea, and Th+Ho+Ea) exhibited the highest significant number of taxa compared with BS (Table S3).

Differences between BS and Ea were more prominent in the upper layer than in the lower layer. In the lower layer, the enriched taxa included several bacterial phyla (Pseudomonadota, Bacteroidota, Calditrichaeota, Ignavibacteriota, and Chloroflexota) and an archaeon (*Candidatus* Woesearchaeota). Similar bacterial phyla were enriched in the upper layer, along with Actinomycetota and Acidimicrobiia. Alpha and Gammaproteobacteria, such as those belonging to the order Hyphomicrobiales (Rhizobiales), including *Methyloceanibacter* and *Nitratireductor*, were exclusively enriched in Ea. Comparing between BS and Th, the upper sediment layer of Th had higher abundances of several Gammaproteobacteria, especially those in the order Chromatiales, than BS, but no difference was found in the lower layer. However, the comparison between BS and Ho, more differences were greater in the lower layer. The enriched taxa included bacteria from the phyla Pseudomonadota and Acidobacteriota, and an archaeon from the phylum *Ca*. Woesearchaeota.

Mixed seagrass patches showed greater differences than mono-species seagrass patches. In the comparison between BS and Ho+Ea, the lower layer was significantly enriched with Pirellulaceae, such as *Novipirellula*, *Lignipirellula*, *Anatilimnocola*, and *Blastopirellula*, as well as archaea from the phyla *Ca.* Lokiarchaeota, *Ca*. Thermoplasmatota, and *Candidatus* Heimdallarchaeota, while the upper layers mainly featured Anaerolineae (Chloroflexota), Verrucomicrobiota, and *Desulfuromonas*. For BS and Th+Ho, more differences were observed in the lower layer, with an increase in certain archaeal species, such as *Candidatus* Thorarchaeota, and bacterial taxa, such as Deltaproteobacteria. For BS and Th+Ea, several gammaproteobacterial taxa of the order Chromatiales and Thiotrichales were found in the upper layer. In the upper layer of the BS-Th+Ho+Ea sediment, several Gammaproteobacteria, including *Thiotrichaceaebacterium*, Thiohalophilus, and Sedimenticola, were more prevalent and exclusively enriched in this specific layer.

### Functional analysis

The samples exhibited consistent functional profiles, with the dominant pathways being carbohydrate, amino acid, and energy metabolism (Figure 6). Interestingly, when comparing the BS and the pool of all seagrass sediments, functional analysis revealed that seagrass samples were significantly enriched in galactose metabolism, nitrogen metabolism, inositol phosphate metabolism, and flavonoid degradation compared with BS samples. In contrast, BS exhibited enrichment in carotenoid biosynthesis, polycyclic aromatic hydrocarbon degradation, xylene degradation, and unsaturated fatty acid biosynthesis (Figure 7A). When analyzing all sediment types together, seagrass-associated sediments exhibited significant enrichment in pathways related to the synthesis of secondary metabolites, cofactors, amino acids, and energy production, whereas BS samples showed no significant enrichment (Figure 7B).

**Figure 6.**
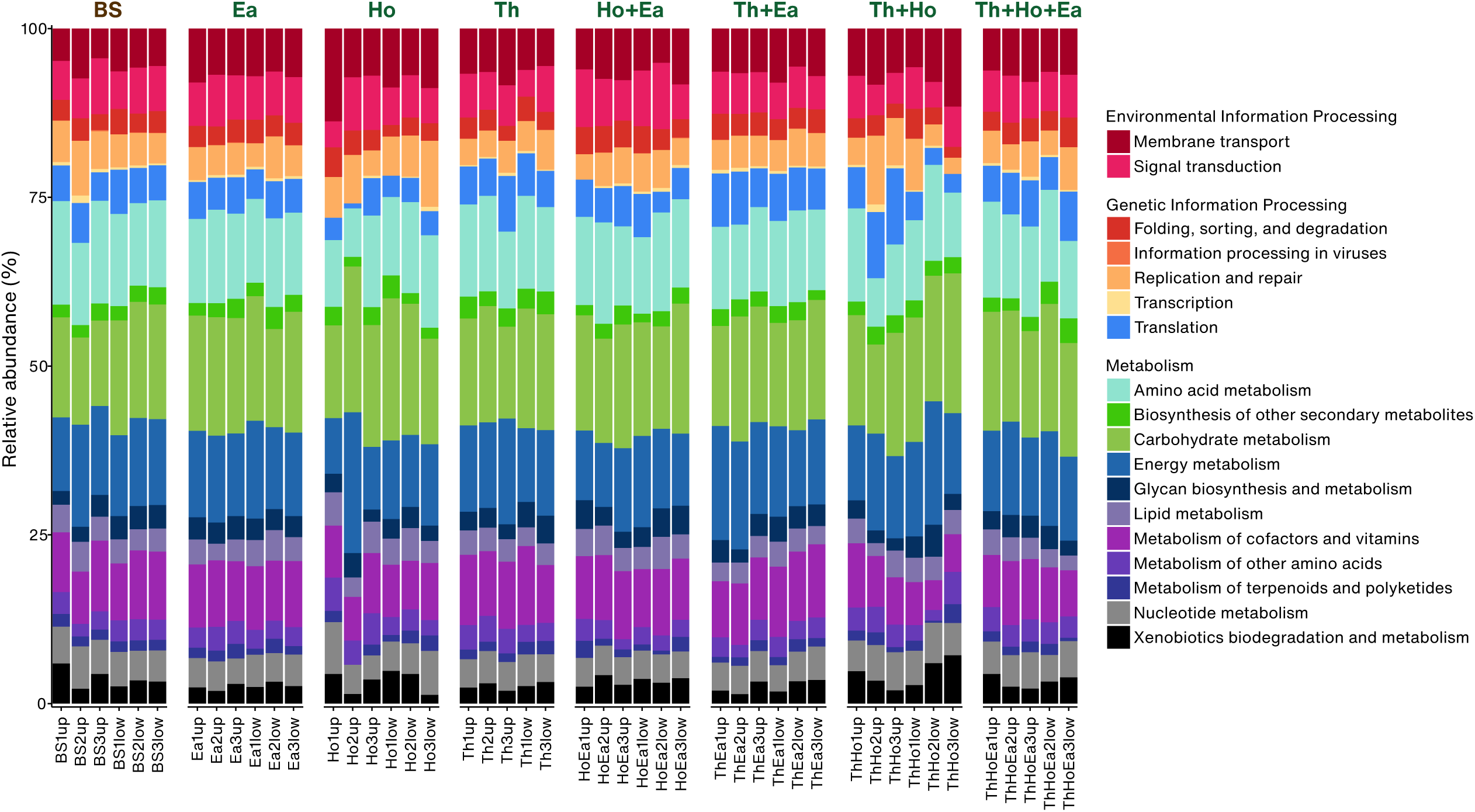
Relative abundance of pathways identified using the Kyoto Encyclopedia of Genes and Genomes database for each sample.

**Figure 7.**
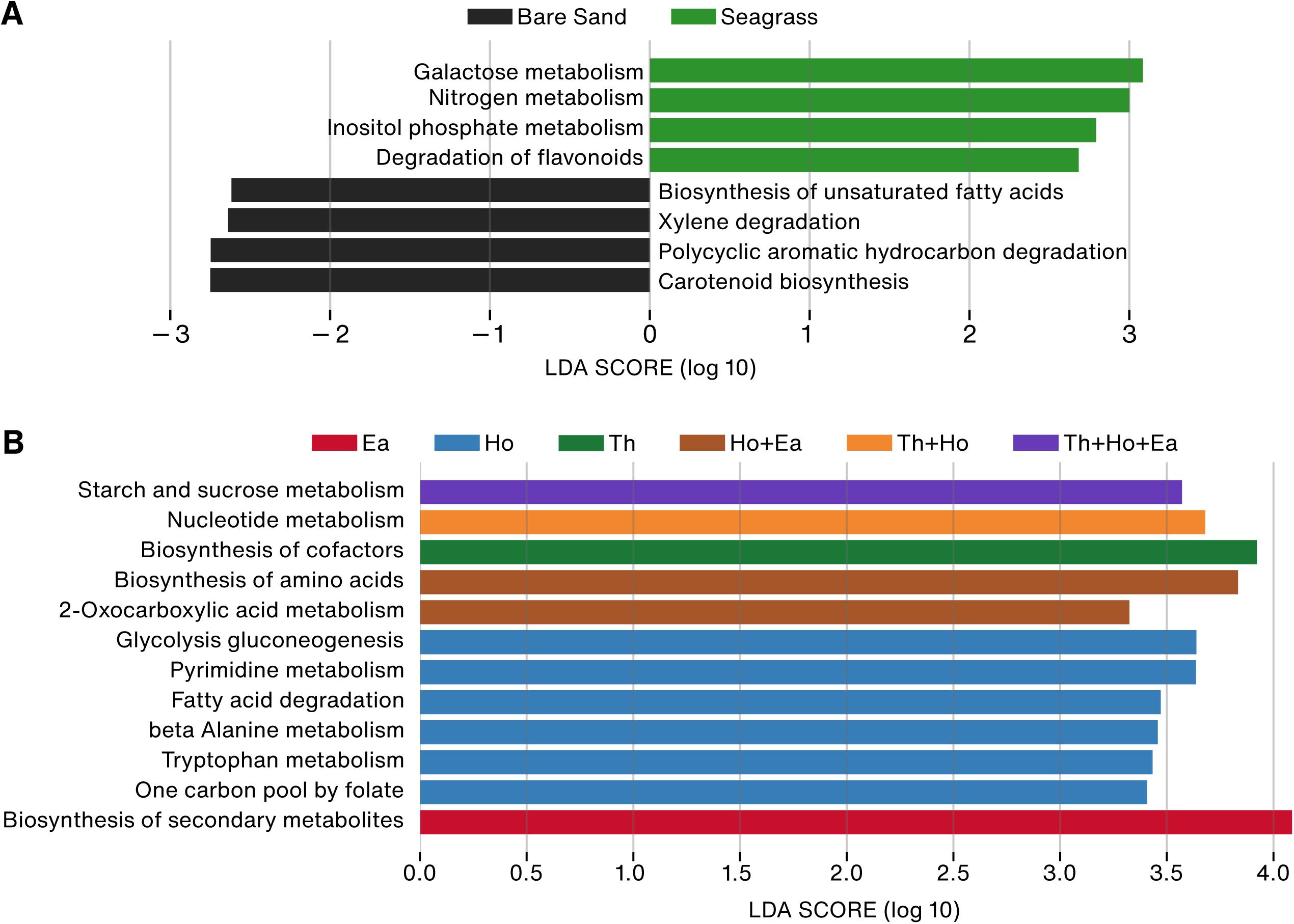
Linear discriminant analysis effect size (LEfSe) analysis of different sediment samples. Comparison between Bare Sand and seagrass sediments (A), and all different types of Bare Sand and seagrass samples (B).

### Metagenome-assembled genomes

Most MAGs belonged to the phyla Pseudomonadota (19 MAGs) and Thermodesulfobacteriota (18 MAGs), while three and one were from the phyla Chloroflexota and Mycoplasmatota, respectively (Table S4). Within the phylum Pseudomonadota, 11 MAGs were placed within *Methyloceanibacter* species (Hyphomicrobiaceae), and two MAGs belonged to a putatively new alphaproteobacteria from the family Rhodospirillaceae (Figure S4A). Five Gammaproteobacteria MAGs of different orders were classified as two new taxa (Figure S4B). Within the phylum Thermodesulfobacteriota, nine MAGs exhibited the closest match to the Desulfobacterales bacterium GCA_016735415.1. Three MAGs were identified as new *Electronema* species, whereas two others were classified under a new taxon belonging to the family Desulfosarcinaceae (Figure S4C). Moreover, the MAGs from the phylum Chloroflexota were classified as members of the Anaerolineales family, and the MAG from the phylum Mycoplasmatota was classified as *Hepatoplasma* (Figure S4D).

Functional analysis of these MAGs revealed that most of them had genes associated with sulfur, hydrogen, and oxygen metabolism. Thermodesulfobacteriota and Pseudomonadota MAGs exhibited similar numbers of genes involved in sulfur metabolism (Table S4). Notably, the dissimilatory sulfite reductase D (*dsrD*) gene was found exclusively in Thermodesulfobacteriota MAGs, whereas the thiosulfate oxidation carrier protein SoxY gene was found solely in Pseudomonadota MAGs (Table S4). Genes associated with hydrogen metabolism were prevalent across most of the analyzed genomes. MAGs belonging to the phylum Thermodesulfobacteriota and class Gammaproteobacteria (phylum Pseudomonadota) had the highest number of genes involved in hydrogen metabolism. Furthermore, only MAGs belonging to the phylum Thermodesulfobacteriota and class Gammaproteobacteria contained genes associated with nitrogen fixation, such as *nifD*, *nifK*, and *nifH*.

## Discussion

### Microbial diversity, substrate, and vertical distribution

Microbial diversity is influenced by various factors, such as sediment depth, water pH, and seagrass species (Banister et al., 2022). We found that the upper sediment layers had a higher alpha diversity of bacteria and archaea, whereas microbial eukaryotes were more abundant in the lower layers. The distribution of some organisms varied across the layers, with some being more abundant in the upper layers and others in the lower layers.

Sediment depth is a key factor in the distribution and diversity of microbes. Among microbial eukaryotes, diatoms (Bacillariophyta) were predominant in both the upper and lower layers, as they are phototrophic organisms that settle on the sediment surface (Alldredge et al., 1995). In the lower layers, their high abundance could be influenced by the reshuffling of the top sediment layer with each tide. Fungi, particularly Ascomycota and Basidiomycota, were more abundance in the lower layer, suggesting their role in the decomposition of organic matter in marine sediments (Moitinho et al., 2022). Our observations regarding the prevalence of archaea are consistent with those of Schorn *et al*. (2022), who also reported a higher abundance of archaea in deeper sediment layers. The decrease in bacterial abundance in the subsurface layers could be due to lower abundance of organic matter, nutrients, and oxygen than the upper sediment layers (Pala et al., 2018; Qiao et al., 2018; Schorn et al., 2022).

Microbial diversity is also influenced by the presence of seagrass species. Because of its unique physical structure, each species can regulate water flow and capture sediment grains of different sizes, altering the sediment’s characteristics and potentially affecting the microbial diversity within seagrass beds (Peterson et al., 2004; Rattanachot and Prathep, 2015). We found that the sediments of *T. hemprichii* had the lowest bacterial and archaeal abundance, contrary to a previous study (Jiang et al., 2015b). We found that *E. acoroides* had the highest number of bacteria, aligning with a study that found that this seagrass species to be effective at trapping waterborne pathogens (Deng et al., 2021). Furthermore, seagrass beds serve as habitats for various organisms and provide shelter for animals, leading to a higher number of eukaryotes residing within them (Bengtsson et al., 2017). This underscores the complex and crucial role of seagrass in their ecosystems.

### Taxonomy

In this study, the predominant organisms in all samples were bacteria, while archaea accounted for around 2.9% of the total taxa. The composition of archaea found in this study is consistent with that shown in previous studies, from less than 2% in the rhizosphere of *Z. marina* on the Atlantic coast of Portugal and France (Cúcio et al., 2018) to 20% in the Mediterranean coastal lagoon (Pala et al., 2018).

The composition of archaea in the seagrass sediments varied among different seagrass species. We found that *Ca*. Bathyarchaeota, Euryarchaeota, and *Ca*. Thermoplasmatota, were the most abundant phyla, consistent with previous studies in marine sediments (P. Liu et al., 2021; Zheng et al., 2019). The phylum Euryarchaeota dominated the upper sediment of all seagrass species. Other studies have shown that seagrass species influence the abundance of archaea. For example, *Ca*. Woesearchaeota was the most abundant phylum in the sediment of *Z. marina* in the Yellow Sea (Zheng et al., 2019), whereas Thermoproteota was predominant in *Z. japonica* meadows (P. Liu et al., 2021). Furthermore, several *Ca*. Woesearchaeota were significantly enriched in Th+Ho+Ea sediments compared with others. *Ca*. Woesearchaeota significantly contributes to nutrient cycles, including anaerobic carbon cycling, nitrogen fixation, and nitrite and sulfate reduction (X. Liu et al., 2021). These findings suggest that seagrass species significantly influence the composition of Archaea in seagrass sediments.

Several sulfur-oxidizing proteobacteria, such as *Thiogranum*, *Thiohalophilus*, and *Electronema*, were abundant in seagrass meadows. They contribute to the nitrogen cycle and are commonly found in vegetated and decomposed sediments (Li et al., 2021; Trevathan-Tackett et al., 2020; Wasmund et al., 2017). They are essential for biological oxidation in coastal marine sediments. Sulfate-reducing and sulfur-oxidizing bacteria were among the most abundant bacteria identified in the seagrass rhizosphere (Cúcio et al., 2016; Fahimipour et al., 2017).

We discovered a substantial number of sequences related to the sulfide-oxidizing bacterial endosymbiont of the lucinid clam *Ctena orbitculata* in the seagrass sediments. Despite the absence of *Ctena* at our study site, we observed another lucinid clam species belonging to the same family, *Rugalucina vietnamica*, commonly distributed in coastal areas. Specifically, *R. vietnamica* inhabits *C. rotundata* beds and bare sand (Rattanachot and Prathep, 2016). The sulfide-oxidizing bacteria endosymbionts of lucinid clams can decrease toxic sulfide levels in seagrass beds. In return, they gain protection from the intricate network of seagrass roots, fostering a mutualistic relationship between the clams and seagrass (Van Der Heide et al., 2012). Moreover, Cúcio *et al*. (2018) reconstructed putative free-living forms of sulfur-oxidizing bacterial symbionts of the clam *Solemya velum* in the seagrass *Z. marina* rhizosphere, suggesting that seagrass meadows could act as a symbiont source for resident invertebrates.

Methane is mainly produced in marine sediments, especially in anoxic environments, by methanogenic archaea via anaerobic degradation of organic matter (Schorn et al., 2022). However, most of the methane is oxidized by microorganisms in the sediment and water column, with a small portion reaching the atmosphere (Reeburgh, 2007). We found that *Methyloceanibacter*, a genus of methane-oxidizing gram-negative bacteria, was more abundant in *E*. *acoroides* sediments. This indicates high methane levels in the sediment, partly because this seagrass species is leafy, produces long roots, and has high biomass (Duarte et al., 1998; Terrados et al., 1998). Previously, a high abundance of *Methyloceanibacter* was reported in decomposing *Zostera muelleri* (Trevathan-Tackett et al., 2020), indicating the use of the methane cycling pathway in these environments (Trevathan-Tackett et al., 2021).

Bacteria involved during the nitrification process were more abundant in BS. Several nitrogen-fixing cyanobacteria of the genus *Crocosphaera* were enriched in BS. These bacteria are widespread in tropical oceans, particularly in areas with high temperatures up to 29 °C (Moisander et al., 2010). This is likely because bare sand tends to have higher temperatures than shaded seagrass patches. Consistant with a previous study in bulk sediment in the Bay of Bengal (Mohapatra et al., 2022), *Nitrospira* (phylum Nitrospirota), nitrite-oxidizing bacteria that prefer low-nitrogen sediments (Nowka et al., 2015), was also significantly enriched in BS.

### Functional analysis

Seagrass sediments are rich in diverse metabolic pathways and microorganisms capable of decomposing various compounds, including carbohydrates such as galactose and inositol phosphates, and flavonoids released by seagrass (Danaraj et al., 2021; Drew, 1983; Suzumura and Kamatani, 1995). The microbes in the sediment use carbon substrates, including carbohydrates, which might vary depending on the seagrass species (Säwström et al., 2016).

The enrichment of the nitrogen metabolism pathway indicates the presence of microorganisms involved in nitrogen cycling, which is important for maintaining nutrient balance and supporting primary productivity in the ecosystem. Bacteria and archaea, including *Phaeobacter* and *Nitrososphaerota*, are essential for phosphorus and nitrogen cycles (Han et al., 2022; Trautwein et al., 2018; Urata et al., 2022; Zhong et al., 2020). This highlights the importance of diverse microbial communities in nutrient cycling and energy flow within the ecosystem.

Pathways involved in the degradation of petroleum products, including xylene and aromatic hydrocarbons were enriched in BS. Several marine bacteria, including *Pseudomonas* and *Rhodococcus*, can degrade xylene (Banerjee et al., 2022). Because these pathways were not observed in sediments associated with seagrass, this emphasizes the importance of seagrass in maintaining the health of marine ecosystems.

### Metagenome-assembled genomes

Desulfobacterales, members of Thermodesulfobacteriota (Desulfobacterota), significantly contribute to the nitrogen and hydrogen cycles in coastal regions (Dyksma et al., 2018; Nie et al., 2021). These are classified under the orders Desulfobacterales and Desulfobulbales, with some identified as new taxa. Among these, several MAGs of sulfate-reducing bacteria could not be assigned to known families.

Among the Pseudomonadota MAGs, four were putatively new Gammaproteobacterial taxa classified under the same order. They possess *nif* genes, indicating their potential roles in nitrogen fixation. Nitrogen-fixing bacteria, which can exist independently or form symbiotic associations with plants, contain *nifD*, *nifK*, and *nifH* genes. For instance, the symbiotic Gammaproteobacteria *Candidatus* Thiodiazotropha of the lucinid clam, a common inhabitant of seagrass beds, have several *nif* genes in their genomes (Osvatic et al., 2021). Interestingly, these genes were not detected in other proteobacterial MAGs.

Belonging to Rhodospirillaceae, two newly discovered alphaproteobacteria MAGs possess genes encoding Rubisco-S and Rubisco-L enzymes. These enzymes are involved in carbon dioxide fixation and are common in alphaproteobacteria (Hügler and Sievert, 2011), but are rarely found in other alphaproteobacterial MAGs. These two MAGs exhibit a low GC content (48%), similar to *Magnetospira* of the same order (Williams et al., 2012). The low GC content suggests that they may be symbionts of other organisms, as bacterial endosymbionts typically have low frequencies of guanine and cytosine (Moran et al., 2009).

### Microbes, seagrass, and warming temperature

Seagrass meadows have been shown to reduce the relative abundance of potential bacterial pathogens that can cause diseases in both humans and marine organisms (Lamb et al., 2017; Tasdemir et al., 2024). The decline of seagrass might prevent the removal of these potential pathogens from the environment.

Seagrass restoration is necessary for carbon storage in the sediments of shallow coastal ecosystems. However, higher seawater temperatures can increase biomass loss for decaying seagrass, change the bacterial community structure, and lower carbon accumulation rates due to increased remineralization rates at the sediment surface. Elevated temperatures negatively affect carbon sequestration rates through enhanced aerobic microbial remineralization, reducing carbon accumulation rates and increasing carbon losses (Trevathan-Tackett et al., 2017). Moreover, increased nutrient load can amplify the contribution of seagrass and algal sources to sediment organic carbon pools, which boosts sediment microbial biomass and extracellular enzyme activity, potentially weakening the seagrass blue carbon sequestration capacity (Liu et al., 2017).

## Conclusion

In conclusion, we found significant differences in several taxa between seagrass and BS sediments and among different seagrass species. The upper parts of the sediments had a higher prevalence of bacteria and microbial eukaryotes, whereas the lower section had a greater abundance of archaea. Seagrass samples were enriched in certain metabolic pathways, such as galactose and flavonoid metabolism, whereas BS showed enrichment in carotenoid and xylene degradation. Moreover, 41 Metagenome-assembled-genomes were recovered, mostly belonging to Pseudomonadota and Thermodesulfobacteriota. However, our study did not comprehensively capture the entire microbial diversity within seagrass environments, indicating the need for future research. Nevertheless, our findings enhance understanding of microbial diversity in seagrass ecosystems, that can facilitate more effective management and restoration strategies for these vital environments.

## Acknowledgments

We thank Piyalap Tuntiprapas and Janmanee Panyawai for helping in sediment collection and Stefano Draisma for commenting on the previous version of the manuscript. The financial support from National Science and Technology Development Agency (NSTDA), Thailand (Code: FDA-CO-2561-7938-TH).

## References

Alldredge, A.L., Gotschalk, C., Passow, U., Riebesell, U., 1995. Mass aggregation of diatom blooms: Insights from a mesocosm study. Deep. Res. Part II Top. Stud. Oceanogr. 42, 9–27. 10.1016/0967-0645(95)00002-8

Alneberg, J., Bjarnason, B.S., De Bruijn, I., Schirmer, M., Quick, J., Ijaz, U.Z., Lahti, L., Loman, N.J., Andersson, A.F., Quince, C., 2014. Binning metagenomic contigs by coverage and composition. Nat. Methods 11, 1144–1146. 10.1038/nmeth.3103

Arkin, A.P., Cottingham, R.W., Henry, C.S., Harris, N.L., Stevens, R.L., Maslov, S., Dehal, P., Ware, D., Perez, F., Canon, S., Sneddon, M.W., Henderson, M.L., Riehl, W.J., Murphy-Olson, D., Chan, S.Y., Kamimura, R.T., Kumari, S., Drake, M.M., Brettin, T.S., Glass, E.M., Chivian, D., Gunter, D., Weston, D.J., Allen, B.H., Baumohl, J., Best, A.A., Bowen, B., Brenner, S.E., Bun, C.C., Chandonia, J.M., Chia, J.M., Colasanti, R., Conrad, N., Davis, J.J., Davison, B.H., Dejongh, M., Devoid, S., Dietrich, E., Dubchak, I., Edirisinghe, J.N., Fang, G., Faria, J.P., Frybarger, P.M., Gerlach, W., Gerstein, M., Greiner, A., Gurtowski, J., Haun, H.L., He, F., Jain, R., Joachimiak, M.P., Keegan, K.P., Kondo, S., Kumar, V., Land, M.L., Meyer, F., Mills, M., Novichkov, P.S., Oh, T., Olsen, G.J., Olson, R., Parrello, B., Pasternak, S., Pearson, E., Poon, S.S., Price, G.A., Ramakrishnan, S., Ranjan, P., Ronald, P.C., Schatz, M.C., Seaver, S.M.D., Shukla, M., Sutormin, R.A., Syed, M.H., Thomason, J., Tintle, N.L., Wang, D., Xia, F., Yoo, H., Yoo, S., Yu, D., 2018. KBase: The United States department of energy systems biology knowledgebase. Nat. Biotechnol. 36, 566–569. 10.1038/nbt.4163

Banerjee, S., Bedics, A., Harkai, P., Kriszt, B., Alpula, N., Táncsics, A., 2022. Evaluating the aerobic xylene-degrading potential of the intrinsic microbial community of a legacy BTEX-contaminated aquifer by enrichment culturing coupled with multi-omics analysis: uncovering the role of Hydrogenophaga strains in xylene degradation. Environ. Sci. Pollut. Res. 29, 28431–28445. 10.1007/s11356-021-18300-w

Banister, R.B., Schwarz, M.T., Fine, M., Ritchie, K.B., Muller, E.M., 2022. Instability and stasis among the microbiome of seagrass leaves, roots and rhizomes, and nearby sediments within a natural pH gradient. Microb. Ecol. 84, 703–716. 10.1007/s00248-021-01867-9

Bengtsson, M.M., Bühler, A., Brauer, A., Dahlke, S., Schubert, H., Blindow, I., 2017. Eelgrass leaf surface microbiomes are locally variable and highly correlated with epibiotic eukaryotes. Front. Microbiol. 8, 1312. 10.3389/fmicb.2017.01312

Brodersen, K.E., Siboni, N., Nielsen, D.A., Pernice, M., Ralph, P.J., Seymour, J., Kühl, M., 2018. Seagrass rhizosphere microenvironment alters plant-associated microbial community composition. Environ. Microbiol. 20, 2854–2864. 10.1111/1462-2920.14245

Celdrán, D., Espinosa, E., Sánchez-Amat, A., Marín, A., 2012. Effects of epibiotic bacteria on leaf growth and epiphytes of the seagrass *Posidonia oceanica*. Mar. Ecol. Prog. Ser. 456, 21–27. 10.3354/meps09672

Chaumeil, P.A., Mussig, A.J., Hugenholtz, P., Parks, D.H., 2020. GTDB-Tk: A toolkit to classify genomes with the genome taxonomy database. Bioinformatics 36, 1925–1927. 10.1093/bioinformatics/btz848

Chen, S., Zhou, Y., Chen, Y., Gu, J., 2018. Fastp: An ultra-fast all-in-one FASTQ preprocessor. Bioinformatics 34, i884–i890. 10.1093/bioinformatics/bty560

Cúcio, C., Engelen, A.H., Costa, R., Muyzer, G., 2016. Rhizosphere microbiomes of European seagrasses are selected by the plant, but are not species specific. Front. Microbiol. 7, 440. 10.3389/fmicb.2016.00440

Cúcio, C., Overmars, L., Engelen, A.H., Muyzer, G., 2018. Metagenomic analysis shows the presence of bacteria related to free-living forms of sulfur-oxidizing chemolithoautotrophic symbionts in the rhizosphere of the seagrass *Zostera marina*. Front. Mar. Sci. 5, 171. 10.3389/fmars.2018.00171

Danaraj, J., Ayyappan, S., Mariasingarayan, Y., Packiyavathy, I.A.S.V., Sweetly Dharmadhas, J., 2021. Chlorophyll fluorescence, dark respiration and metabolomic analysis of *Halodule pinifolia* reveal potential heat responsive metabolites and biochemical pathways under ocean warming. Mar. Environ. Res. 164, 105248. 10.1016/j.marenvres.2020.105248

Deng, Y., Liu, S., Feng, J., Wu, Y., Mao, C., 2021. What drives putative bacterial pathogens removal within seagrass meadows? Mar. Pollut. Bull. 166, 112229. 10.1016/j.marpolbul.2021.112229

Drew, E.A., 1983. Sugars, cytolitols and seagrass phylogeny. Aquat. Bot. 15, 387–408. 10.1016/0304-3770(83)90007-4

Duarte, C.M., Merino, M., Agawin, N.S.R., Uri, J., Fortes, M.D., Gallegos, M.E., Marbá, N., Hemminga, M.A., 1998. Root production and belowground seagrass biomass. Mar. Ecol. Prog. Ser. 171, 97–108. 10.3354/meps171097

Dyksma, S., Pjevac, P., Ovanesov, K., Mussmann, M., 2018. Evidence for H2 consumption by uncultured Desulfobacterales in coastal sediments. Environ. Microbiol. 20, 450–461. 10.1111/1462-2920.13880

Fahimipour, A.K., Kardish, M.R., Lang, J.M., Green, J.L., Eisen, J.A., Stachowicz, J.J., 2017. Global-scale structure of the eelgrass microbiome. Appl. Environ. Microbiol. 83, e03391–16. 10.1128/AEM.03391-16

Fortes, M.D., Ooi, J.L.S., Tan, Y.M., Prathep, A., Bujang, J.S., Yaakub, S.M., 2018. Seagrass in Southeast Asia: A review of status and knowledge gaps, and a road map for conservation. Bot. Mar. 61, 269–288. 10.1515/bot-2018-0008

Han, Y., Zhang, M., Chen, X., Zhai, W., Tan, E., Tang, K., 2022. Transcriptomic evidences for microbial carbon and nitrogen cycles in the deoxygenated seawaters of Bohai Sea. Environ. Int. 158, 106889. 10.1016/j.envint.2021.106889

Hügler, M., Sievert, S.M., 2011. Beyond the Calvin cycle: Autotrophic carbon fixation in the ocean. Ann. Rev. Mar. Sci. 3, 261–289. 10.1146/annurev-marine-120709-142712

Jiang, Y.F., Ling, J., Dong, J. De, Chen, B., Zhang, Y.Y., Zhang, Y.Z., Wang, Y.S., 2015a. Illumina-based analysis the microbial diversity associated with *Thalassia hemprichii* in Xincun Bay, South China Sea. Ecotoxicology 24, 1548–1556. 10.1007/s10646-015-1511-z

Jiang, Y.F., Ling, J., Wang, Y.S., Chen, B., Zhang, Y.Y., Dong, J. De, 2015b. Cultivation-dependent analysis of the microbial diversity associated with the seagrass meadows in Xincun Bay, South China Sea. Ecotoxicology 24, 1540–1547. 10.1007/s10646-015-1519-4

Kanehisa, M., Sato, Y., Morishima, K., 2016. BlastKOALA and GhostKOALA: KEGG tools for functional characterization of genome and metagenome sequences. J. Mol. Biol. 428, 726–731. 10.1016/j.jmb.2015.11.006

Kang, D.D., Li, F., Kirton, E., Thomas, A., Egan, R., An, H., Wang, Z., 2019. MetaBAT 2: An adaptive binning algorithm for robust and efficient genome reconstruction from metagenome assemblies. PeerJ 2019, e7359. 10.7717/peerj.7359

Lamb, J.B., Van De Water, J.A.J.M., Bourne, D.G., Altier, C., Hein, M.Y., Fiorenza, E.A., Abu, N., Jompa, J., Harvell, C.D., 2017. Seagrass ecosystems reduce exposure to bacterial pathogens of humans, fishes, and invertebrates. Science (80-. ). 355, 731–733. 10.1126/science.aal1956

Li, M., Fang, A., Yu, X., Zhang, K., He, Z., Wang, C., Peng, Y., Xiao, F., Yang, T., Zhang, W., Zheng, X., Zhong, Q., Liu, X., Yan, Q., 2021. Microbially-driven sulfur cycling microbial communities in different mangrove sediments. Chemosphere 273, 128597. 10.1016/j.chemosphere.2020.128597

Liu, P., Zhang, H., Song, Z., Huang, Y., Hu, X., 2021. Seasonal dynamics of Bathyarchaeota-dominated benthic archaeal communities associated with seagrass (*Zostera japonica*) meadows. J. Mar. Sci. Eng. 9, 1304. 10.3390/JMSE9111304

Liu, S., Jiang, Z., Zhang, J., Wu, Y., Huang, X., Macreadie, P.I., 2017. Sediment microbes mediate the impact of nutrient loading on blue carbon sequestration by mixed seagrass meadows. Sci. Total Environ. 599–600, 1479–1484. 10.1016/j.scitotenv.2017.05.129

Liu, S., Wu, Y., Luo, H., Ren, Y., Jiang, Z., Zhang, X., Fang, Y., Liang, J., Huang, X., 2023. Seagrass canopy structure mediates putative bacterial pathogen removal potential. Front. Mar. Sci. 9, 1076097. 10.3389/fmars.2022.1076097

Liu, X., Wang, Y., Gu, J.D., 2021. Ecological distribution and potential roles of Woesearchaeota in anaerobic biogeochemical cycling unveiled by genomic analysis. Comput. Struct. Biotechnol. J. 19, 794–800. 10.1016/j.csbj.2021.01.013

Lyimo, L.D., Gullström, M., Lyimo, T.J., Deyanova, D., Dahl, M., Hamisi, M.I., Björk, M., 2018. Shading and simulated grazing increase the sulphide pool and methane emission in a tropical seagrass meadow. Mar. Pollut. Bull. 134, 89–93. 10.1016/j.marpolbul.2017.09.005

Martin, B.C., Alarcon, M.S., Gleeson, D., Middleton, J.A., Fraser, M.W., Ryan, M.H., Holmer, M., Kendrick, G.A., Kilminster, K., 2020. Root microbiomes as indicators of seagrass health. FEMS Microbiol. Ecol. 96, fiz201. 10.1093/femsec/fiz201

McLeod, E., Chmura, G.L., Bouillon, S., Salm, R., Björk, M., Duarte, C.M., Lovelock, C.E., Schlesinger, W.H., Silliman, B.R., 2011. A blueprint for blue carbon: Toward an improved understanding of the role of vegetated coastal habitats in sequestering CO2. Front. Ecol. Environ. 9, 552–560. 10.1890/110004

McMurdie, P.J., Holmes, S., 2013. Phyloseq: An R Package for reproducible interactive analysis and graphics of microbiome census data. PLoS One 8, e61217. 10.1371/journal.pone.0061217

Menzel, P., Ng, K.L., Krogh, A., 2016. Fast and sensitive taxonomic classification for metagenomics with Kaiju. Nat. Commun. 7, 11257. 10.1038/ncomms11257

Mikheenko, A., Prjibelski, A., Saveliev, V., Antipov, D., Gurevich, A., 2018. Versatile genome assembly evaluation with QUAST-LG. Bioinformatics 34, i142–i150. 10.1093/bioinformatics/bty266

Mohapatra, M., Manu, S., Dash, S.P., Rastogi, G., 2022. Seagrasses and local environment control the bacterial community structure and carbon substrate utilization in brackish sediments. J. Environ. Manage. 314, 115013. 10.1016/j.jenvman.2022.115013

Moisander, P.H., Beinart, R.A., Hewson, I., White, A.E., Johnson, K.S., Carlson, C.A., Montoya, J.P., Zehr, J.P., 2010. Unicellular cyanobacterial distributions broaden the oceanic N2 fixation domain. Science (80-. ). 327, 1512–1514. 10.1126/science.1185468

Moitinho, M.A., Chiaramonte, J.B., Bononi, L., Gumiere, T., Melo, I.S., Taketani, R.G., 2022. Fungal succession on the decomposition of three plant species from a Brazilian mangrove. Sci. Rep. 12, 14547. 10.1038/s41598-022-18667-x

Moran, N.A., McLaughlin, H.J., Sorek, R., 2009. The dynamics and time scale of ongoing genomic erosion in symbiotic bacteria. Science (80-. ). 323, 379–382. 10.1126/science.1167140

Mou, X., Sun, S., Edwards, R.A., Hodson, R.E., Moran, M.A., 2008. Bacterial carbon processing by generalist species in the coastal ocean. Nature 451, 708–711. 10.1038/nature06513

Nie, S., Zhang, Z., Mo, S., Li, J., He, S., Kashif, M., Liang, Z., Shen, P., Yan, B., Jiang, C., 2021. Desulfobacterales stimulates nitrate reduction in the mangrove ecosystem of a subtropical gulf. Sci. Total Environ. 769, 144562. 10.1016/j.scitotenv.2020.144562

Nowka, B., Daims, H., Spieck, E., 2015. Comparison of oxidation kinetics of nitrite-oxidizing bacteria: Nitrite availability as a key factor in niche differentiation. Appl. Environ. Microbiol. 81, 745–753. 10.1128/AEM.02734-14

Osvatic, J.T., Wilkins, L.G.E., Leibrecht, L., Leray, M., Zauner, S., Polzin, J., Camacho, Y., Gros, O., van Gils, J.A., Eisen, J.A., Petersen, J.M., Yuen, B., 2021. Global biogeography of chemosynthetic symbionts reveals both localized and globally distributed symbiont groups. Proc. Natl. Acad. Sci. U. S. A. 118, e2104378118. 10.1073/pnas.2104378118

Pala, C., Molari, M., Nizzoli, D., Bartoli, M., Viaroli, P., Manini, E., 2018. Environmental drivers controlling bacterial and archaeal abundance in the sediments of a Mediterranean lagoon ecosystem. Curr. Microbiol. 75, 1147–1155. 10.1007/s00284-018-1503-3

Parks, D.H., Imelfort, M., Skennerton, C.T., Hugenholtz, P., Tyson, G.W., 2015. CheckM: Assessing the quality of microbial genomes recovered from isolates, single cells, and metagenomes. Genome Res. 25, 1043–1055. 10.1101/gr.186072.114

Peterson, C.H., Luettich, R.A., Micheli, F., Skilleter, G.A., 2004. Attenuation of water flow inside seagrass canopies of differing structure. Mar. Ecol. Prog. Ser. 268, 81–92. 10.3354/meps268081

Prjibelski, A., Antipov, D., Meleshko, D., Lapidus, A., Korobeynikov, A., 2020. Using SPAdes De Novo Assembler. Curr. Protoc. Bioinforma. 70, e102. 10.1002/cpbi.102

Qiao, Y., Liu, J., Zhao, M., Zhang, X.H., 2018. Sediment depth-dependent spatial variations of bacterial communities in mud deposits of the Eastern China marginal seas. Front. Microbiol. 9, 1128. 10.3389/fmicb.2018.01128

Rattanachot, E., Prathep, A., 2016. The effect of increasing seagrass root complexity and redox potential on the population of *Pillucina vietnamica* (Bivalvia: Lucinidae) in southwestern Thailand. Molluscan Res. 36, 142–151. 10.1080/13235818.2015.1128587

Rattanachot, E., Prathep, A., 2015. Species specific effects of three morphologically different belowground seagrasses on sediment properties. Estuar. Coast. Shelf Sci. 167, 427–435. 10.1016/j.ecss.2015.10.019

Reeburgh, W.S., 2007. Oceanic methane biogeochemistry. Chem. Rev. 107, 486–513. 10.1021/cr050362v

Säwström, C., Serrano, O., Rozaimi, M., Lavery, P.S., 2016. Utilization of carbon substrates by heterotrophic bacteria through vertical sediment profiles in coastal and estuarine seagrass meadows. Environ. Microbiol. Rep. 8, 582–589. 10.1111/1758-2229.12406

Schorn, S., Ahmerkamp, S., Bullock, E., Weber, M., Lott, C., Liebeke, M., Lavik, G., Kuypers, M.M.M., Graf, J.S., Milucka, J., 2022. Diverse methylotrophic methanogenic archaea cause high methane emissions from seagrass meadows. Proc. Natl. Acad. Sci. U. S. A. 119, e2106628119. 10.1073/pnas.2106628119

Seemann, T., 2014. Prokka: Rapid prokaryotic genome annotation. Bioinformatics 30, 2068–2069. 10.1093/bioinformatics/btu153

Segata, N., Izard, J., Waldron, L., Gevers, D., Miropolsky, L., Garrett, W.S., Huttenhower, C., 2011. Metagenomic biomarker discovery and explanation. Genome Biol. 12, R60. 10.1186/gb-2011-12-6-r60

Shaffer, M., Borton, M.A., McGivern, B.B., Zayed, A.A., La Rosa, S.L. 0003 3527 8101, Solden, L.M., Liu, P., Narrowe, A.B., Rodríguez-Ramos, J., Bolduc, B., Gazitúa, M.C., Daly, R.A., Smith, G.J., Vik, D.R., Pope, P.B., Sullivan, M.B., Roux, S., Wrighton, K.C., 2020. DRAM for distilling microbial metabolism to automate the curation of microbiome function. Nucleic Acids Res. 48, 8883–8900. 10.1093/nar/gkaa621

Sieber, C.M.K., Probst, A.J., Sharrar, A., Thomas, B.C., Hess, M., Tringe, S.G., Banfield, J.F., 2018. Recovery of genomes from metagenomes via a dereplication, aggregation and scoring strategy. Nat. Microbiol. 3, 836–843. 10.1038/s41564-018-0171-1

Stankovic, M., Ambo-Rappe, R., Carly, F., Dangan-Galon, F., Fortes, M.D., Hossain, M.S., Kiswara, W., Van Luong, C., Minh-Thu, P., Mishra, A.K., Noiraksar, T., Nurdin, N., Panyawai, J., Rattanachot, E., Rozaimi, M., Soe Htun, U., Prathep, A., 2021. Quantification of blue carbon in seagrass ecosystems of Southeast Asia and their potential for climate change mitigation. Sci. Total Environ. 783, 146858. 10.1016/j.scitotenv.2021.146858

Suzumura, M., Kamatani, A., 1995. Mineralization of inositol hexaphosphate in aerobic and anaerobic marine sediments: Implications for the phosphorus cycle. Geochim. Cosmochim. Acta 59, 1021–1026. 10.1016/0016-7037(95)00006-2

Tasdemir, D., Scarpato, S., Utermann-Thüsing, C., Jensen, T., Blümel, M., Wenzel-Storjohann, A., Welsch, C., Echelmeyer, V.A., 2024. Epiphytic and endophytic microbiome of the seagrass *Zostera marina*: Do they contribute to pathogen reduction in seawater? Sci. Total Environ. 908, 168422. 10.1016/j.scitotenv.2023.168422

Terrados, J., Duarte, C.M., Fortes, M.D., Borum, J., Agawin, N.S.R., Bach, S., Thampanya, U., Kamp-Nielsen, L., Kenworthy, W.J., Geertz-Hansen, O., Vermaat, J., 1998. Changes in community structure and biomass of seagrass communities along gradients of siltation in SE Asia. Estuar. Coast. Shelf Sci. 46, 757–768. 10.1006/ecss.1997.0304

Titioatchasai, J., Surachat, K., Rattanachot, E., Tuntiprapas, P., Mayakun, J., 2023. Assessment of diversity of marine organisms among natural and transplanted seagrass meadows. J. Mar. Sci. Eng. 11, 1928. 10.3390/jmse11101928

Trautwein, K., Hensler, M., Wiegmann, K., Skorubskaya, E., Wöhlbrand, L., Wünsch, D., Hinrichs, C., Feenders, C., Müller, C., Schell, K., Ruppersberg, H., Vagts, J., Koßmehl, S., Steinbüchel, A., Schmidt-Kopplin, P., Wilkes, H., Hillebrand, H., Blasius, B., Schomburg, D., Rabus, R., 2018. The marine bacterium *Phaeobacter inhibens* secures external ammonium by rapid buildup of intracellular nitrogen stocks. FEMS Microbiol. Ecol. 94, fiy154. 10.1093/femsec/fiy154

Trevathan-Tackett, S.M., Jeffries, T.C., Macreadie, P.I., Manojlovic, B., Ralph, P., 2020. Long-term decomposition captures key steps in microbial breakdown of seagrass litter. Sci. Total Environ. 705, 135806. 10.1016/j.scitotenv.2019.135806

Trevathan-Tackett, S.M., Kepfer-Rojas, S., Engelen, A.H., York, P.H., Ola, A., Li, J., Kelleway, J.J., Jinks, K.I., Jackson, E.L., Adame, M.F., Pendall, E., Lovelock, C.E., Connolly, R.M., Watson, A., Visby, I., Trethowan, A., Taylor, B., Roberts, T.N.B., Petch, J., Farrington, L., Djukic, I., Macreadie, P.I., 2021. Ecosystem type drives tea litter decomposition and associated prokaryotic microbiome communities in freshwater and coastal wetlands at a continental scale. Sci. Total Environ. 782, 146819. 10.1016/j.scitotenv.2021.146819

Trevathan-Tackett, S.M., Seymour, J.R., Nielsen, D.A., Macreadie, P.I., Jeffries, T.C., Sanderman, J., Baldock, J., Howes, J.M., Steven, A.D.L., Ralph, P.J., 2017. Sediment anoxia limits microbial-driven seagrass carbon remineralization under warming conditions. FEMS Microbiol. Ecol. 93, fix033. 10.1093/femsec/fix033

Urata, S., Kurosawa, Y., Yamasaki, N., Yamamoto, H., Nishiwaki, N., Hongo, Y., Adachi, M., Yamaguchi, H., 2022. Utilization of phosphonic acid compounds by marine bacteria of the genera *Phaeobacter*, *Ruegeria*, and *Thalassospira* (α-Proteobacteria). FEMS Microbiol. Lett. 369, fnac065. 10.1093/femsle/fnac065

Van Der Heide, T., Govers, L.L., De Fouw, J., Olff, H., Van Der Geest, M., Van Katwijk, M.M., Piersma, T., Van De Koppel, J., Silliman, B.R., Smolders, A.J.P., Van Gils, J.A., 2012. A three-stage symbiosis forms the foundation of seagrass ecosystems. Science (80-. ). 336, 1432–1434. 10.1126/science.1219973

Wasmund, K., Mußmann, M., Loy, A., 2017. The life sulfuric: microbial ecology of sulfur cycling in marine sediments. Environ. Microbiol. Rep. 9, 323–344. 10.1111/1758-2229.12538

Williams, T.J., Lefèvre, C.T., Zhao, W., Beveridge, T.J., Bazylinski, D.A., 2012. *Magnetospira thiophila* gen. nov., sp. nov., a marine magnetotactic bacterium that represents a novel lineage within the Rhodospirillaceae (Alphaproteobacteria). Int. J. Syst. Evol. Microbiol. 62, 2443–2450. 10.1099/ijs.0.037697-0

Wu, Y.W., Simmons, B.A., Singer, S.W., 2016. MaxBin 2.0: An automated binning algorithm to recover genomes from multiple metagenomic datasets. Bioinformatics 32, 605–607. 10.1093/bioinformatics/btv638

Zheng, P., Wang, C., Zhang, X., Gong, J., 2019. Community structure and abundance of Archaea in a *Zostera marina* meadow: A comparison between seagrass-colonized and bare sediment sites. Archaea 2019, 5108012. 10.1155/2019/5108012

Zhong, H., Zhong, H., Lehtovirta-Morley, L., Liu, J., Liu, J., Zheng, Y., Lin, H., Song, D., Todd, J.D., Tian, J., Zhang, X.H., Zhang, X.H., Zhang, X.H., 2020. Novel insights into the Thaumarchaeota in the deepest oceans: Their metabolism and potential adaptation mechanisms. Microbiome 8, 78. 10.1186/s40168-020-00849-2

