## Supplementary material for "Metagenomics insights into microbial diversity shifts among seagrass sediments": Figure S1

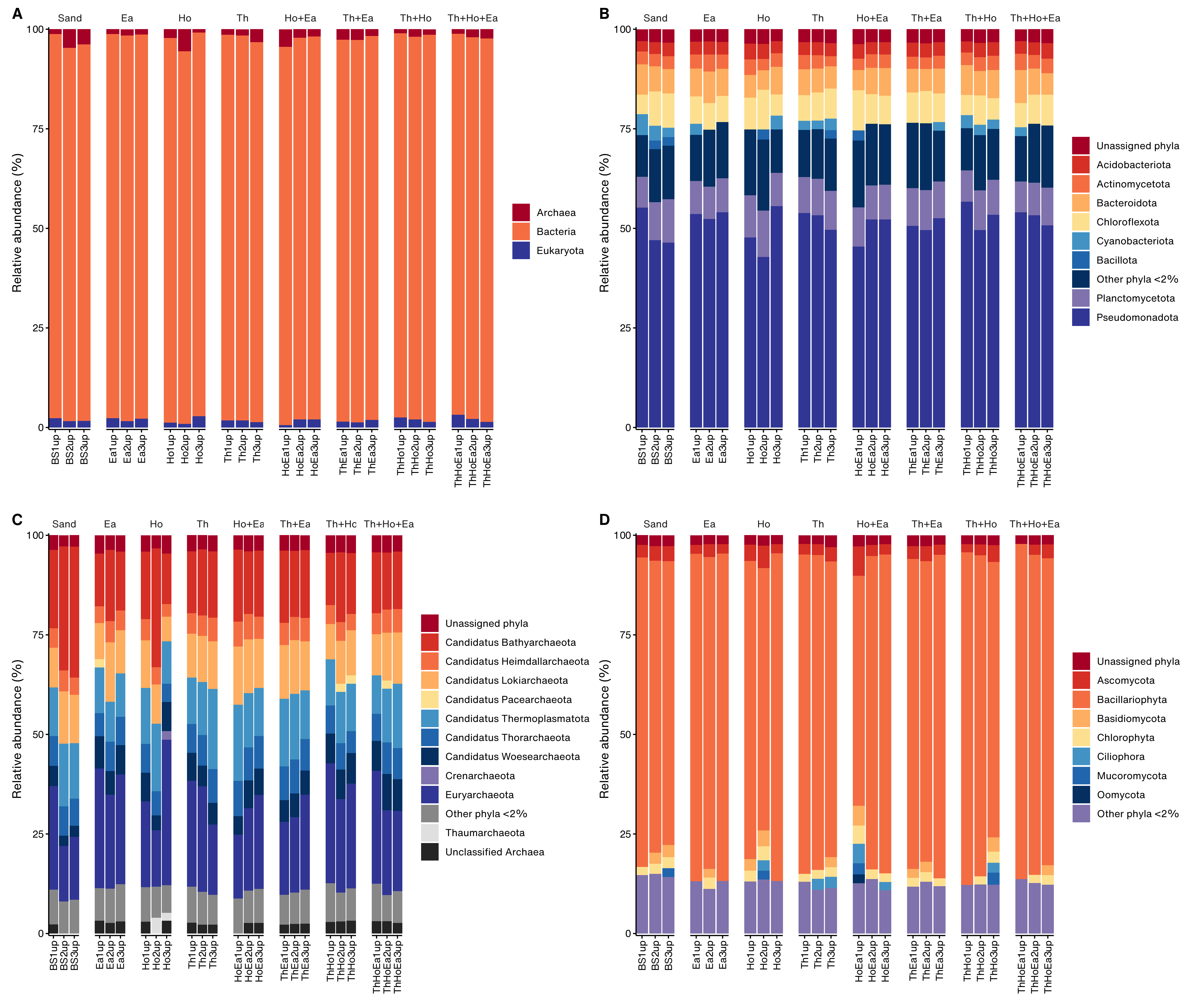

**Figure S1.** Bar plots represent relative abundance of cellular organisms in the upper layers of the sediments in each sample. At the superkingdom level (A), the phylum level of bacteria (B), archaea (C), and microbial eukaryotes (D).

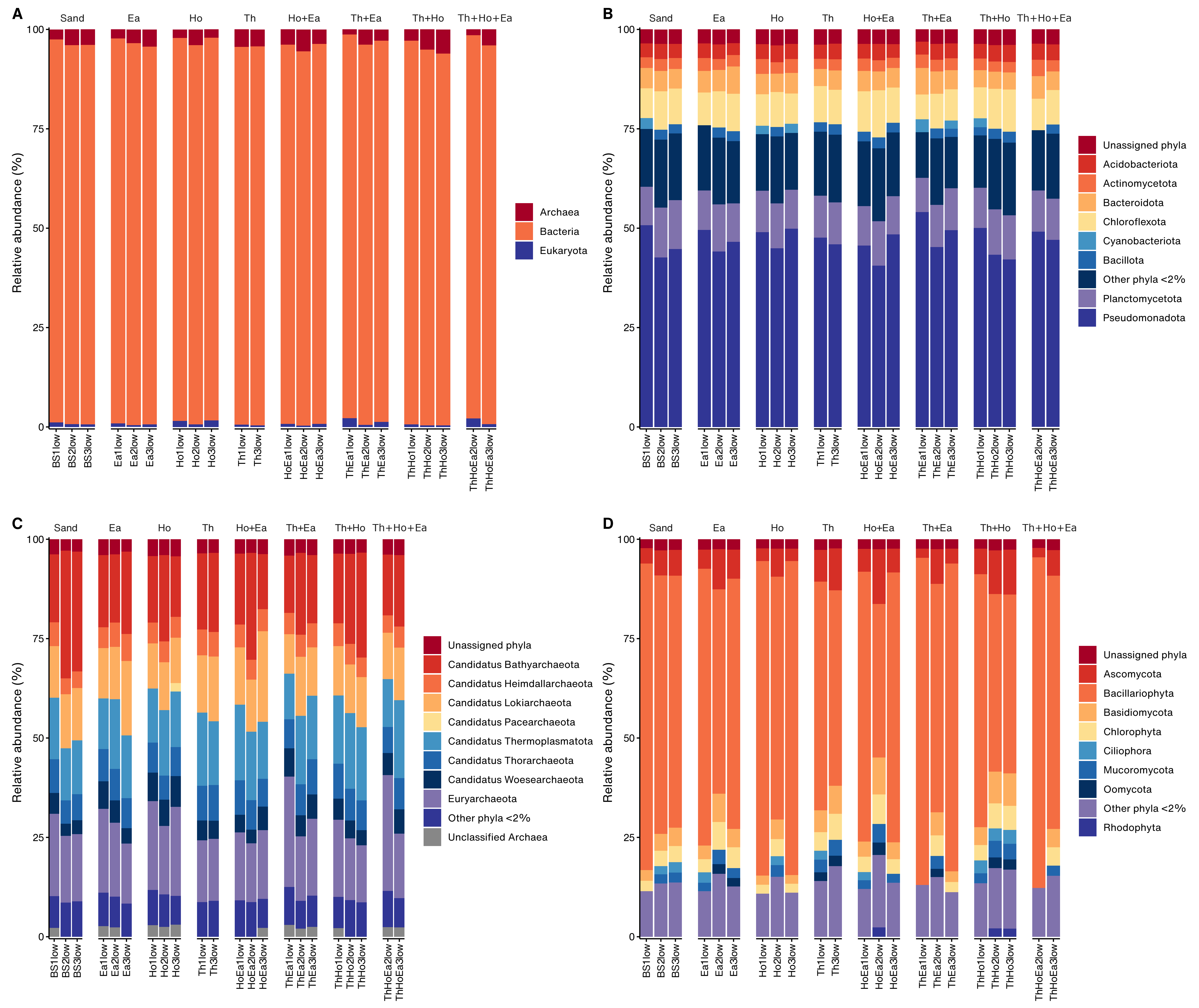

**Figure S2.** Bar plots represent relative abundance of cellular organisms in the lower layers of the sediments in each sample. At the superkingdom level (A), the phylum level of bacteria (B), archaea (C), and microbial eukaryotes (D).

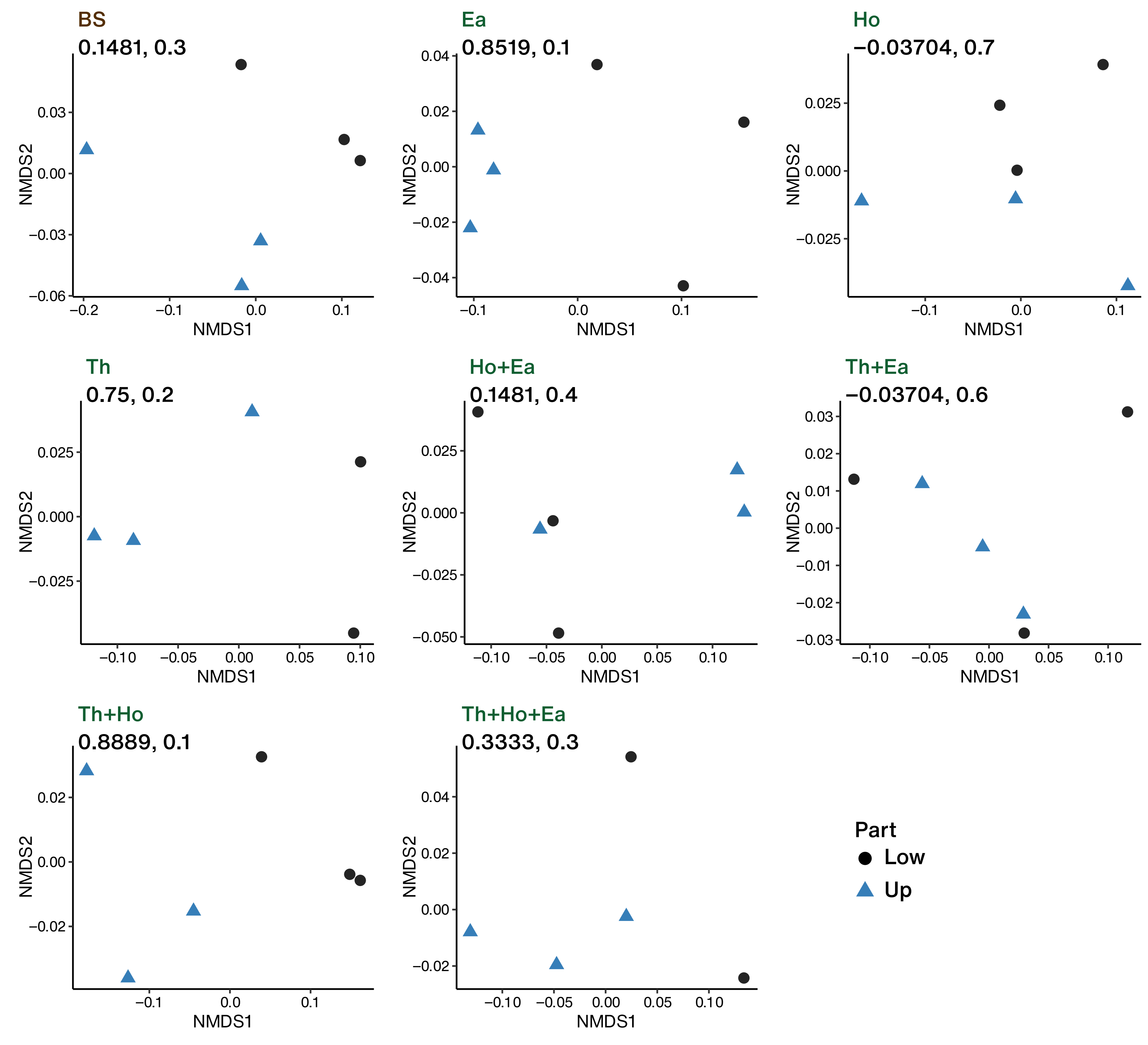

**Figure S3.** Nonmetric multidimensional scaling based on the Bray-Curtis distance of each sample. Values indicate ANOSIM statistic R and p-value between the upper and the lower layers.

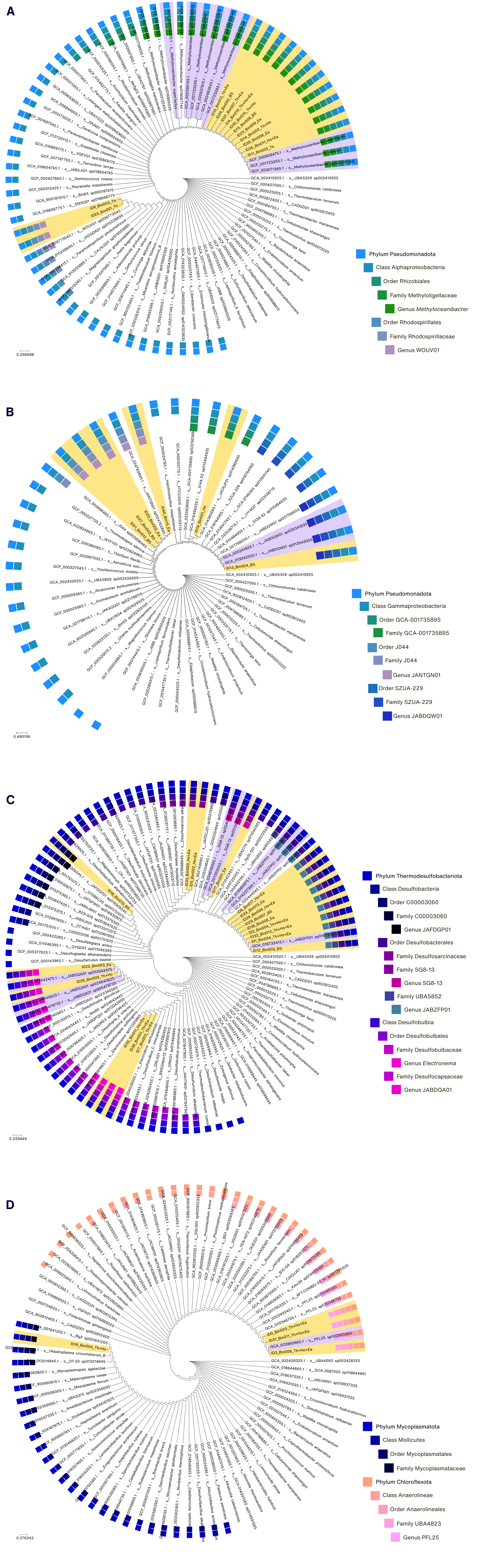

**Figure S4.** Phylogenetic trees of metagenome-assembled genomes (MAGs) reconstructed in this study. (A) and (B) Phylum Pseudomonadota. (C) Phylum Thermodesulfobacteriota. (D) Phylum Mycoplasmatota and Chloroflexota. Yellow labels indicate MAGs found in this study.
